# Staining and resin embedding of whole *Daphnia* magna samples for micro-CT imaging enabling 3D visualization of cells, tissues, and organs of various thicknesses

**DOI:** 10.1101/2023.05.21.541654

**Authors:** Mee S. Ngu, Daniel J. Vanselow, Rachelle A. Saint-Fort, Andrew L. Sugarman, Carolyn R. Zaino, Maksim A. Yakovlev, Keith C. Cheng, Khai C. Ang

## Abstract

Micro-CT imaging is a powerful tool for generating high resolution, isotropic three-dimensional datasets of whole, centimeter-scale model organisms that can be used for qualitative and quantitative analysis. The small size, global freshwater distribution, wide range of cell size and structures of micron scale, and common use of *D. magna* in toxicological and environmental studies make it an ideal model for demonstrating the potential power of micro-CT-enabled whole-organism phenotyping. This protocol details the steps involved in *D. magna* samples preparation for micro-CT: euthanasia, fixation, staining, and resin embedding. Micro-CT reconstructions of samples imaged using synchrotron micro-CT reveal histological (microanatomic) features of organ systems, tissues, and cells in the context of the entire organism at sub-micron resolution, and in 3 dimensionality. The enabled “3D histology” and 3D renderings can be used towards morphometric analyses across cells, tissues, and organ systems for both descriptive and hypothesis testing studies.

## Introduction

Micro-computed tomography (micro-CT) is increasingly recognized as a valuable imaging technique for generating three-dimensional datasets that enable 3D visualization, and qualitative and quantitative analysis of biological samples. Imaging of whole, intact samples allows detailed investigation of overall morphology and cellular structures is especially useful for evaluation of microanatomy and phenotypes in various model organisms [1–5]. *Daphnia magna* is a keystone branchiopod crustacean (order Cladocera) in freshwater lotic ecosystems worldwide and is an established model in ecology and evolution [6,7]. They have a parthenogenetic life cycle that allows the rearing of identical clones’ populations from single genotypes. Their short generation time also permits the experimental manipulation of large populations for concurrent study of molecular and phenotypic responses to stressors. They are responsive to environmental change, and has been commonly used for evaluating environmental stressors [8–11] and toxicity testing [12–15]. Studies using this crustacean model for the characterization of abnormalities and pathological change in whole animal can greatly benefit from micro-CT imaging that allows 3D examination at sub-micron voxel resolution [16].

Sample preparation that results in undistorted and well-preserved microanatomical features is the first step in the generation of high-quality micro-CT images. This protocol details the sample preparation of whole *D. magna* for micro-CT imaging including euthanasia, fixation, staining with metal, and resin embedding. Metal staining is useful for micro-CT imaging because the inherent contrast between different soft tissues in micro-CT images is weak. Phosphotungstic acid (PTA), a heteropoly acid with the chemical formula H_3_PWO_40_, is one of the most widely used contrast agents for micro-CT imaging because it provides superior contrast between tissue components [17]. However, PTA staining of invertebrates can take up to one week or longer [18–21]. The commonly used PTA concentration of 0.3% works for staining small juveniles (instar 1-3 or 1-3 days after extruding from brood chamber) within 48 hours but does not provide uniform staining for gravid adults (instar 8 and older, S1 Fig.). We show here that a higher concentration of PTA (3%) can be used to provide homogenous staining of the whole *D. magna* adults in 3 days. Samples can be kept in ethanol if it can be scanned immediately. If immediate imaging is not needed or available, sample dehydration and embedding in resin [22] is recommended because samples stored in ethanol deteriorate over time. The scope and scale of images made possible by the protocol provided is a necessary step towards enabling computational morphological analysis of genetic and environmental change in *Daphnia* and other Cladocera species.

## Materials and methods

The detailed protocol described here is included with this article as Step-by-step protocol (S1 File) and will be published in protocol.io.

### *Daphnia magna* culturing

A commercial strain of *D. magna* was purchased from Carolina Biological (NC, USA) and raised in “Aachener Daphnien-Medium” or ADaM at room temperature (20°C ± 1°C) under a 16-hour light/8-hour dark photoperiod. *D. magna* cultures were fed three times weekly with 3.0 x 10^7^ cells/ml of green microalgae (*Raphidocelis subcapitata)* and once a week with 0.1 mg/mL of dissolved bakers’ yeast.

### Euthanasia, Fixation and Staining

*D. magna* samples were euthanized in carbonated water and fixed in Bouin’s solution for 24 h before staining in PTA for at least 48 h depending on size or age of the samples. Serial-dehydration using 70%, 90%, 95% and 100% ethanol, followed by resin-embedding of samples in LR White were recommended for both immediate imaging or long-term storage of samples and data re-acquisition.

### Micro-CT Imaging and Reconstruction

Scans were performed using a custom benchtop system and synchrotron system at beamline 8.3.2 at the Advanced Light Source at Lawrence Berkeley National Laboratory. Synchrotron scans were acquired using a 5 mm field-of-view/0.5 um pixel resolution imaging system at 20 keV, as a sequence of 150 ms projections [16]. Depending on the diameter of samples, about 3000 projections were obtained over 180° for adult females. Additionally, 20 flat-field (gain) images (at the beginning and end of acquisition) and 20 dark-field images were also acquired. Flat-field correction, stripe removal, and image reconstruction were performed using the open source TomoPy toolkit [23]. Reconstructions resulted in isotropic voxel size of 0.52 µm^3^.

Custom benchtop micro-CT system images were collected using an Indium Gallium liquid metal jet X-ray source (Excillum D2+) with a LuAG (Metal-laser) scintillator and a 10mm field-of-view/0.7 system. Source anode voltage was set to 70kV and 150W. 500 projections were taken of each sample. Exposure time per projection was 1200ms. Samples were rotated with continuous motion over 220 degrees during each imaging session. Source to sample distance was 208 mm. To avoid the use of cone beam reconstruction, source to scintillator distance was 19 mm. The camera (Vieworks VP-151MC) was set to hardware SUM bin4 to boost signal.

Reconstructions were performed using parallel geometry with the gridrec algorithm in Tomopy [23]. Final image volumes achieved a voxel size of 2.8 µm^3^.

## Results

The protocol described here details sample preparation of whole *Daphnia* for micro-CT imaging. Resulting 3D reconstructions reveal anatomic (organ) and micro-anatomic (cellular) features in the context of the entire organism. Micro-CT reconstructions from our custom-built benchtop scanner at 2.8 µm voxel resolution allow the visualization of various organs (S2 Fig). Synchrotron-based micro-CT reconstructions at 0.5 µm resolution adds cellular details within each organ of the whole organism scans. The similarity of digital slices to conventional histology output is demonstrated by representative coronal, sagittal and transverse cross-sections (Fig 1). Highlights of some cellular components include cellular details in the optic lobe and cerebrum ganglia, features in the ovary (yolks, lipid droplets, nurse cells and nucleus of an egg), fat cell nucleoli (about 10 µm in diameter), gut epithelial cell nucleoli (about 2.5 µm in diameter), and cellular details in the developing embryos (Fig 2). Individual intestinal microvilli cannot be visualized because they are smaller than 1 micron in diameter. However, their collective directionality, perpendicular to the gut surface, is represented by the texture of the brush border (Fig 2D).

**Fig 1.**
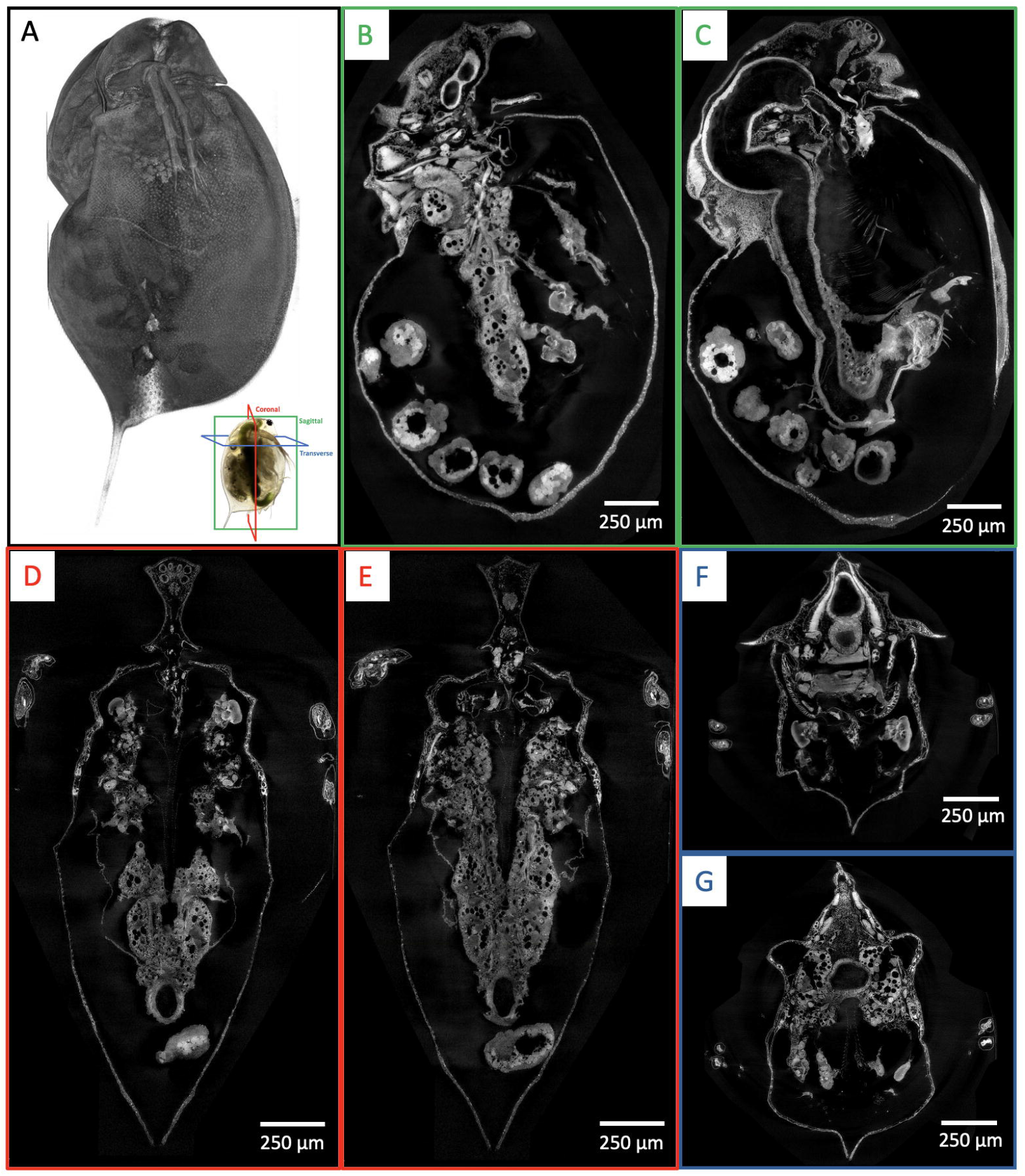
Whole organism imaging of PTA-stained *D. magna* at cell resolution enables histology-like cross sections. 3D volume rendering (A) shows the scanning electron microscopy-like surface rendering. Sagittal (B, C), coronal (D, E) and transverse (F and G) cross sections can be obtained from one sample after imaging. B-G represent 5 μm thick micro-CT slabs.

**Fig 2.**
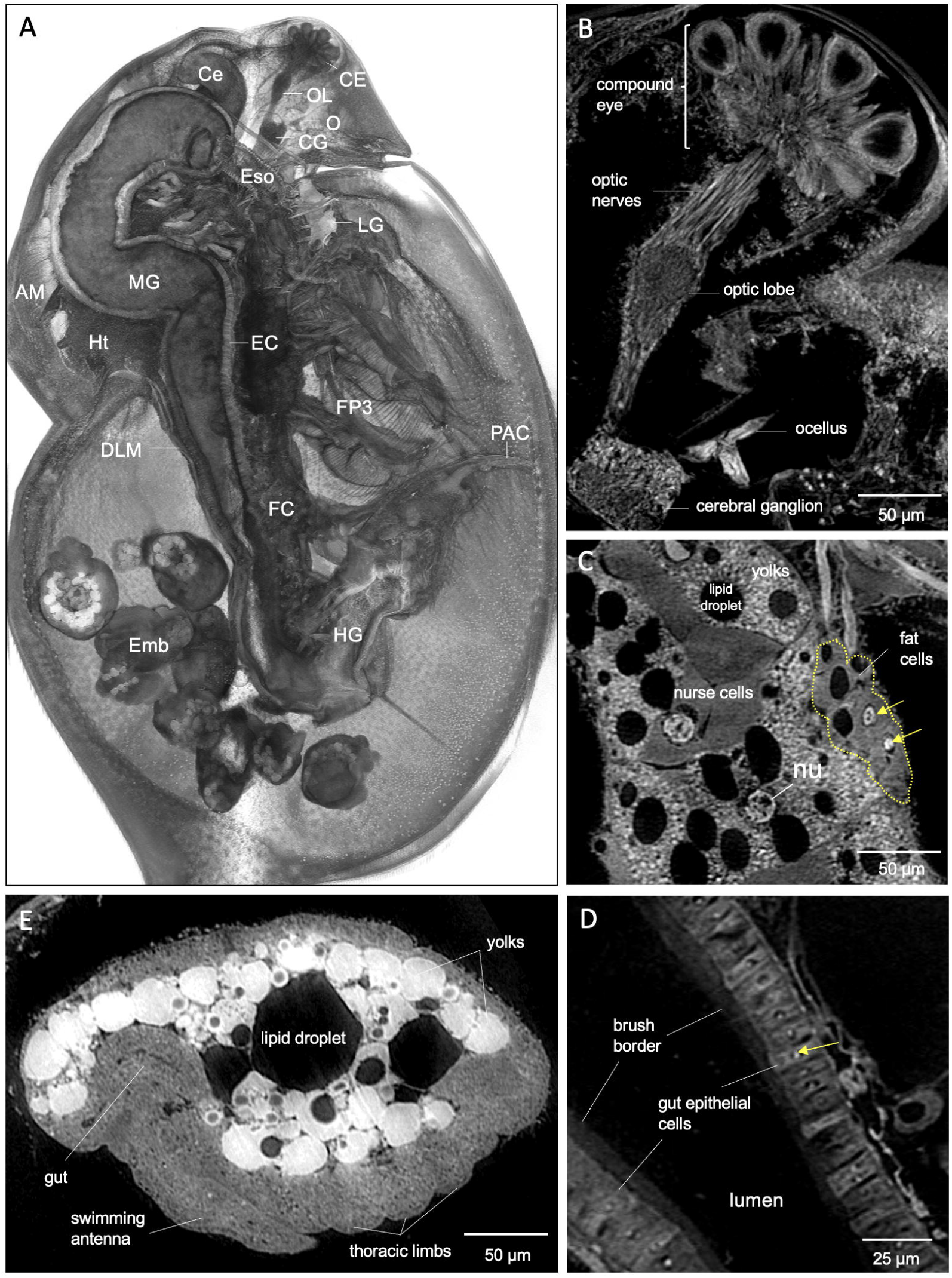
Microanatomic features of an adult female *D. magna* from synchrotron-based micro-CT imaging at 0.5 μm per pixel resolution. (A) 3D rendering at mid-section of the sagittal plane with various organs, organ substructures, and cell types indicated. AM, antennal muscles; Ce, hepatic ceca; CE, compound eye; CG, cerebral ganglia; DLM, dorsal longitudinal muscles; EC, gut epithelial cells, Emb, developing embryos; Eso, esophagus; FC, fat cells; FP3; filter plates on third pair of thoracic limbs; HG, hindgut; Ht, heart; LG, labral glands; O, ocellus, OL; optic lobe; PAC, post-abdomen claws. Highlights of microanatomical features are such as: (B) Cellular details and connections between the compound eye, optic nerves, optic lobe, cerebral ganglia, and ocellus. (C) Details in the ovary showing nucleus (nu) of the oocyte, nurse cells, yolks, and lipid droplets. Nucleoli (yellow arrows) in fat cells are also clearly visible. (D) Nucleoli (yellow arrows) in the gut epithelial cells, and brush border. (E) Recognizable details in the developing embryo include precursors of the gut, swimming antennae and thoracic limbs. B and C represent 5 μm thick micro-CT slabs; (D) and (E) represent individual 0.5 μm thick micro-CT slices.

While both traditional histology and micro-CT imaging have the resolution needed to distinguish cellular features in 2D slice, only the latter can reveal thicker, complex 3D tissue structures such as heart and paired filter plates on the third and fourth pairs of thoracic limbs. Customizing micro-CT slab thicknesses for specific anatomical structures enable visualization of whole organs in details (Fig 3). Slabs are generated from maximum intensity projection of micro-CT slices. Besides slabs, visualization of multiple anatomical structures such as the compound eye, optic nerves, optic lobe, cerebral ganglia, and ocellus in the vision system, can also be customized using 3D rendering with various viewing angles and cutting planes (Fig 4).

**Fig 3.**
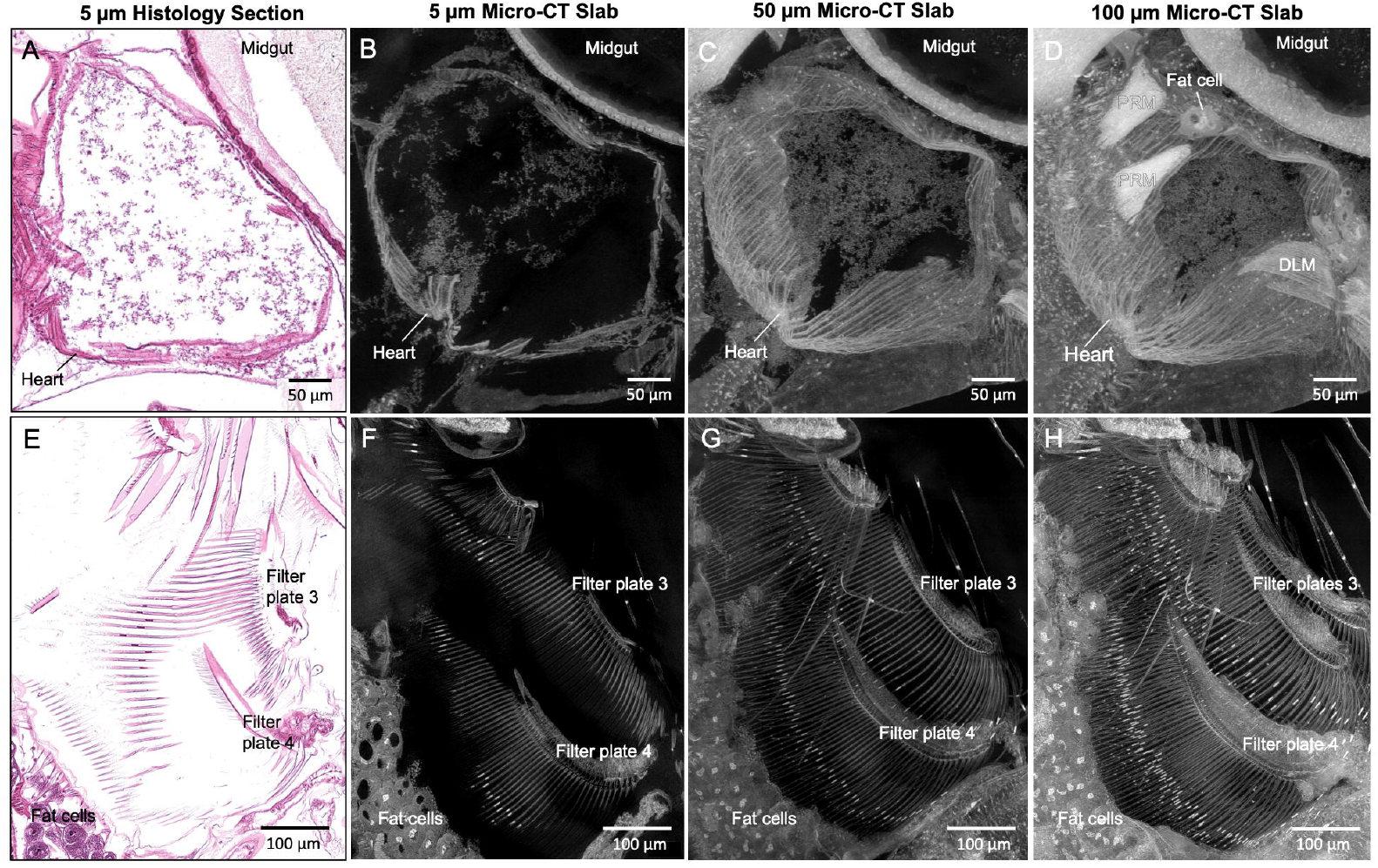
Visualization of anatomical structures using micro-CT slabs of various thicknesses. 5 μm thick micro-CT slab of heart and filter plates (B and F, respectively) resembling the 5 μm thick histological tissue section (A and E, respectively). Thicker micro-CT slabs (50 μm) allow visualization of more heart wall muscles (C) and long setae of the filter plates (G). Micro-CT slabs of 100 μm showed the posterior rotator muscles of mandible (PRM) and fat cells around the heart (D), and both filter plates on thoracic limb 3 (H). DLM, dorsal longitudinal muscle.

**Fig 4.**
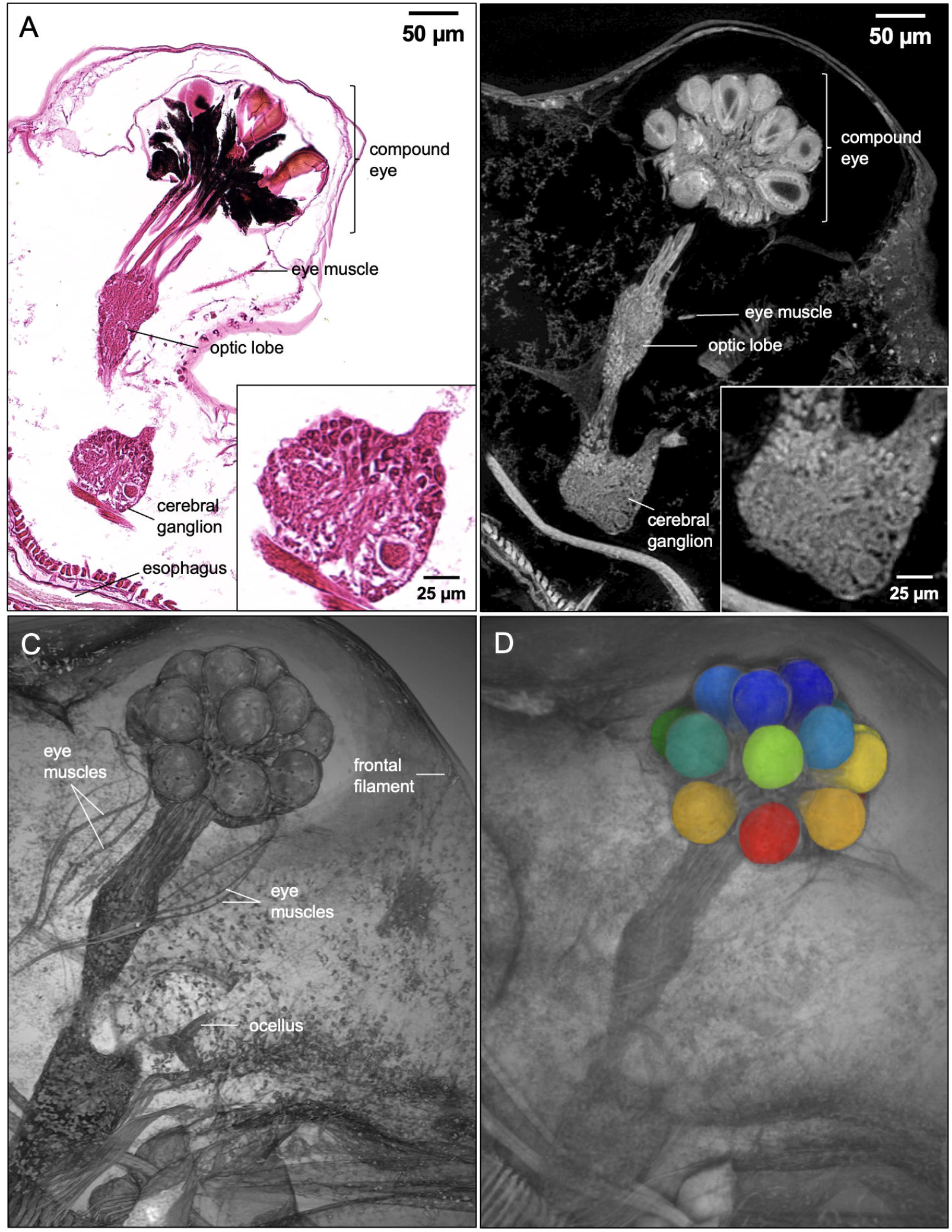
Histology section, micro-CT images and 3D rendering of *D. magna* vision system. Comparison of *D. magna* visual system using 5 µm thick histology section and micro-CT slab of the same thickness (A and B, respectively). Insets show detail of cerebral ganglion where the micro-CT slab demonstrates the near histological resolution of micro-CT imaging at 0.5-micron resolution. (C) 3D rendering featuring the vision system allows the clear visibility of the frontal filament that is connected to ocellus, the eye muscle bundles, and their insertion to the compound eye. (D) Image segmentation of structures of interested (crystalline cones shown here) allows the isolation of specific structures for measurement or quantitative analysis.

## Discussion

In developing this protocol, we prioritized time-efficiency and sufficient detail for newcomers to prepare whole *Daphnia* samples for micro-CT imaging. Bouin’s is the fixative of choice for whole *Daphnia* samples because paraformaldehyde and 10% neutral buffered formalin yield less consistent fixation. Samples fixed overnight in paraformaldehyde and 10% neutral buffered formalin tend to exhibit a fixation artifact in which the carapace is expanded, and the post-abdomen is extended ventrally, causing embryos present to be dislodged from the brood chamber [24]. Fixation of several arthropod taxa in Bouin’s has also been reported to provide better results in terms of tissue contrast when compared with ethanol and glutaraldehyde solution [25].

PTA stain is commonly used at concentrations of 0.3-0.5% for micro-CT imaging [17,22]. While the lower concentration of 0.3% PTA is sufficient for small/young *D. magna* juveniles, it does not provide homogenous staining of gravid adults after 72 hours. In contrast, 3% PTA provides homogenous staining of gravid adult samples in 72 hours resulting in ideal contrast for high-resolution micro-CT imaging. For adult samples carrying many developing embryos (>15), an additional 24 hours is needed to ensure that all the embryos are stained completely. Renewal of PTA solution after 48 hours of incubation is important for achieving homogenous staining. The optimal PTA concentrations and time efficient staining durations to achieve even contrast for samples of various ages is summarized in **Table 1**.

Resin-embedded *Daphnia* samples are suitable for both immediate imaging and long-term storage of samples and data re-acquisition. Sample preparation involving critical-point drying [25] and drying by chemical (hexamethyldisilazane) [26] is possible. However, solid supportive matrix can prevent appendage movement during imaging that will result in motion artifact.

The above protocol designed for micro-CT imaging of *Daphnia* is applicable to other Cladocera and may be adaptable to other chitinous terrestrial invertebrates of similar size for broader taxonomic, ecological, anatomic, genetic, and toxicological studies.

## Supporting Information

### S1 File. Step-by-step protocol

**S1 Fig. Uneven staining of adult gravid *D. magna***. Only some muscles and epipodites of the thoracic limbs, portion of the developing embryos and carapace were stained in 0.3% PTA after 48h.

**S2 Fig. Anatomic features of an adult female shown by scan from benchtop micro-CT scanner at 2.8** μ**m per pixel resolution**. AM, antennal muscles; Ce, hepatic ceca; CE, compound eye; CG, cerebral ganglia; DLM, dorsal longitudinal muscles; Emb, developing embryos; Ht, heart; LG, labral glands; O, ocellus, OL; optic lobe.

## Acknowledgements

The authors are grateful to Dr. Dilworth Parkinson and Advanced Light Sources Beamline 8.3.2, Lawrence Berkeley National Labs for making possible micro-CT imaging of the *Daphnia* samples.

