## Supplementary material for "Staining and resin embedding of whole *Daphnia* magna samples for micro-CT imaging enabling 3D visualization of cells, tissues, and organs of various thicknesses": S1 File

**S1 File. Step-by-step protocol**

All liquid volumes used for incubation, rinse, and infiltration represented at least 20X sample volume. All incubation, rinse, and infiltration steps are performed on a low-speed orbital shaker (Corning LSE) set to 55 revolutions per minute (RPM) to encourage agitation without damaging the samples by rubbing against the container surface.

1. Fixation

    1. Transfer the *Daphnia* sample using bulb pipette containing the least amount of water into a flat bottom glass vial such as a 20ml scintillator vial (Wheaton) filled with carbonated water to euthanize the *Daphnia*. As soon as the sample stops moving, transfer it into Bouin's solution. (Note: Tip of bulb pipette trimmed at a 45° angle such that the diameter is bigger than the size of the sample).
    2. Immediately, replace the Bouin's solution and fix the sample in fresh Bouin's solution at room temperature for 24 hours (hrs). (Note: replacing with fresh Bouin's is to remove any water carried over)

2. Staining

    1. Rinse the fixed sample with 1x phosphate buffered saline (PBS) pH 7.4 for 10 minutes (min), thrice.
    2. Submerge the sample in 35% ethyl alcohol (EtOH) for 15 min at room temperature with gentle agitation.
    3. Discard the 35% EtOH and submerge the sample in 50% EtOH for 15 min at room temperature with gentle agitation.

4. Discard the 50% EtOH and submerge sample in PTA (in 70% EtOH) at room temperature with gentle agitation. Concentration of PTA and staining duration depend on the sample's age (Table 1). Replacement of PTA stain solution is highly recommended after 48 hrs, especially for samples with developing embryos in the brood chamber. Pre-made PTA should be stirred for 15 min before use to prevent deposition of PTA that can appear as random bright spots in samples in reconstruction.

Table 1. Concentration of PTA and staining duration for different ages of *D. magna*

| Age | Embryos in brood chamber | PTA concentration | Staining duration |
|---|---|---|---|
| Juvenile (instar 1-3) | no | 0.3% | 48 hrs |
| Juvenile (instar 4-7) | no | 1% | 48 hrs |
| Adult (instar 8 and older) | yes | 3% | 72 hrs or longer (with a PTA renewal at 48 hrs) |

5. After staining, wash the sample in 70% EtOH for 5 min with gentle agitation, twice.
6. For adult sample to be scanned in 70% EtOH, transfer the sample using a trimmed bulb pipette or 1 ml pipette tip into a 200 μl micropipette tip and seal the tip end with polymer oven-bake clay (Sculpey). For smaller juvenile samples, 10 μl micropipette tips can be used.
7. Tap the sealed end gently to release bubbles.
8. Use clean round tip forceps or wooden stick to GENTLY push the sample down the pipette tip until it is immobilized against the wall of the pipette tip, without squeezing or crushing it.

9. Seal the opening of the pipette tip with parafilm to avoid evaporation and the sample is ready to be scanned (Fig 1). In general, PTA-stained samples can be stored up to a month. If samples do not need to be permanent, scanning *Daphnia* samples at this step is possible within 3 days without ballooning artifact. To avoid ballooning artifact and to allow imaging of *Daphnia* more than 3 days after preparation we recommend serial dehydration and resin-embedding (below).

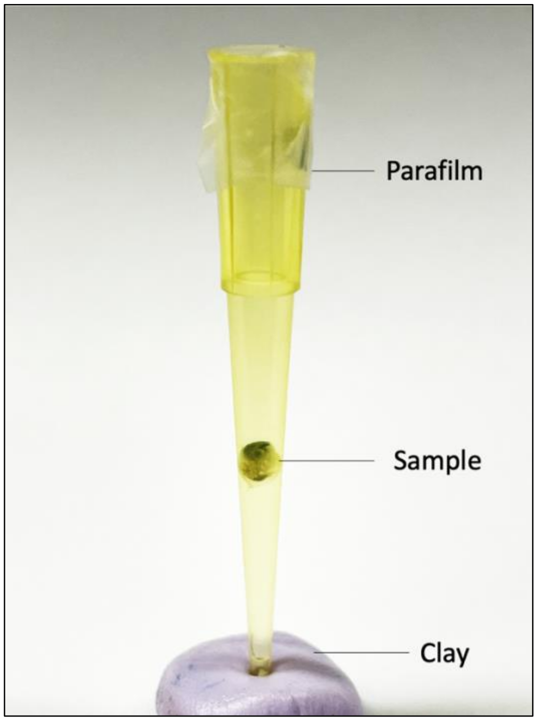


**Fig 1. Gravid *D. magna* sample in 200 µl micropipette tip for imaging.**

3. Dehydration and Resin Embedding (optional). Dehydration and embedding of samples in LR White acrylic resin allows long-term storage and reacquisition, which is convenient when immediate scanning is not possible.

1. For dehydration, submerge the sample in 90%, 95%, 100% and 100% EtOH for 20 min each concentration at room temperature with gentle agitation.
2. Prepare 1:1 v/v mixture of 100% EtOH and LR White acrylic resin. Submerge the samples in 1:1 EtOH and LR White acrylic resin mixture overnight or at least 3 hrs at room temperature with gentle agitation.
3. Submerge the sample in 100% LR White resin for 2 hrs at room temperature with gentle agitation.
4. Replace the LR White resin with fresh 100% LR White resin and submerge for 1 hr at room temperature with gentle agitation.
5. Cut a polyimide tubing of appropriate diameter to 30 - 50 mm length (Table 2).

Table 2. Sizes of polyimide tubing and micropipette tip for different ages of *D. magna*

| Age | Polyimide tubing inner diameter | Micropipette tip |
|---|---|---|
| Juvenile (instar 1-3) | 0.0403” | 200 µl (end of tip clipped off) |
| Juvenile (instar 4-7) | 0.0808” | 200 µl |
| Adult (instar 8 and older) | 0.105” | 1000 µl |

6. Attach the polyimide tubing to a micropipette tip so that it fits snugly (Fig 2). (Note: 200 µl micropipette tip for 0.0403” tubing and 1000 µl micropipette tip for 0.105” tubing. Clip off the end of 200 µl micropipette tip for the tubing to fit snugly.)

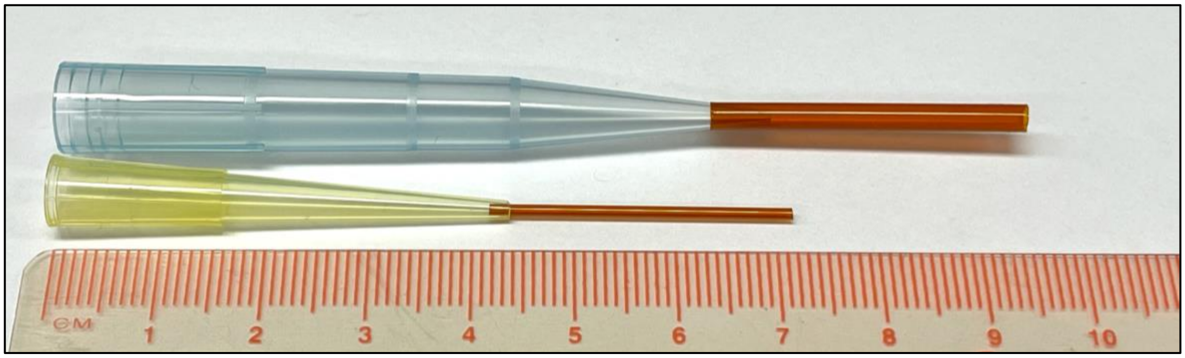


**Fig 2. Polyimide tubing attached to micropipette tips for LR White resin embedding.** The end of 200 μl micropipette tip was clipped off at 4.4 cm for 0.0403” polyimide tubing to fit snugly. 0.105” polyimide tubing can be attached snugly to the 1000 μl micropipette tip without clipping off the end.

7. Attach the micropipette tip together with the polyimide tubing to a micropipette.
8. Transfer the sample to a small weigh boat or V-shaped solution basin and fully submerge the samples in fresh 100% LR White resin.
9. Position the tubing at the head of the sample and pipette resin to fill up half of the tubing before pipetting the specimen slowly into the tubing. (Pipette the sample into the tubing such that the sample is moving in the natural, forward direction to avoid backward movement that will damage extremities). Position the sample in the middle of the tubing and ensure the tubing above and below the specimen is filled with resin.
10. Immediately seal the open end tightly, using a slab of oven-bake clay that has been flattened into a ~2mm-thick sheet. Remove excess clay.
11. Separate the micropipette tip from the polyimide tube by gentle rotation and seal that end of the tube with clay.

12. Place the tubing horizontally, with one end slightly elevated to minimize bubble formation next to the sample. Allow the resin to polymerize for 24 hrs at 65 °C. Once polymerized, the sample is ready for imaging (Fig  3).

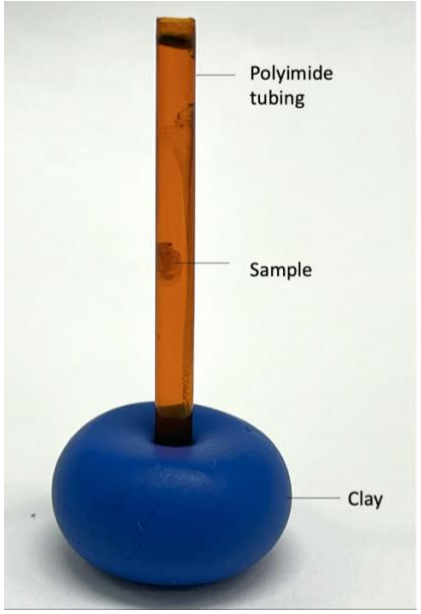


**Fig  3. Sample embedded in LR White acrylic resin using polyimide tubing.** Removal of polyimide tubing is possible but not necessary for imaging.
