## Supplementary figures and images for "Staining and resin embedding of whole *Daphnia* magna samples for micro-CT imaging enabling 3D visualization of cells, tissues, and organs of various thicknesses"

### S1 Fig

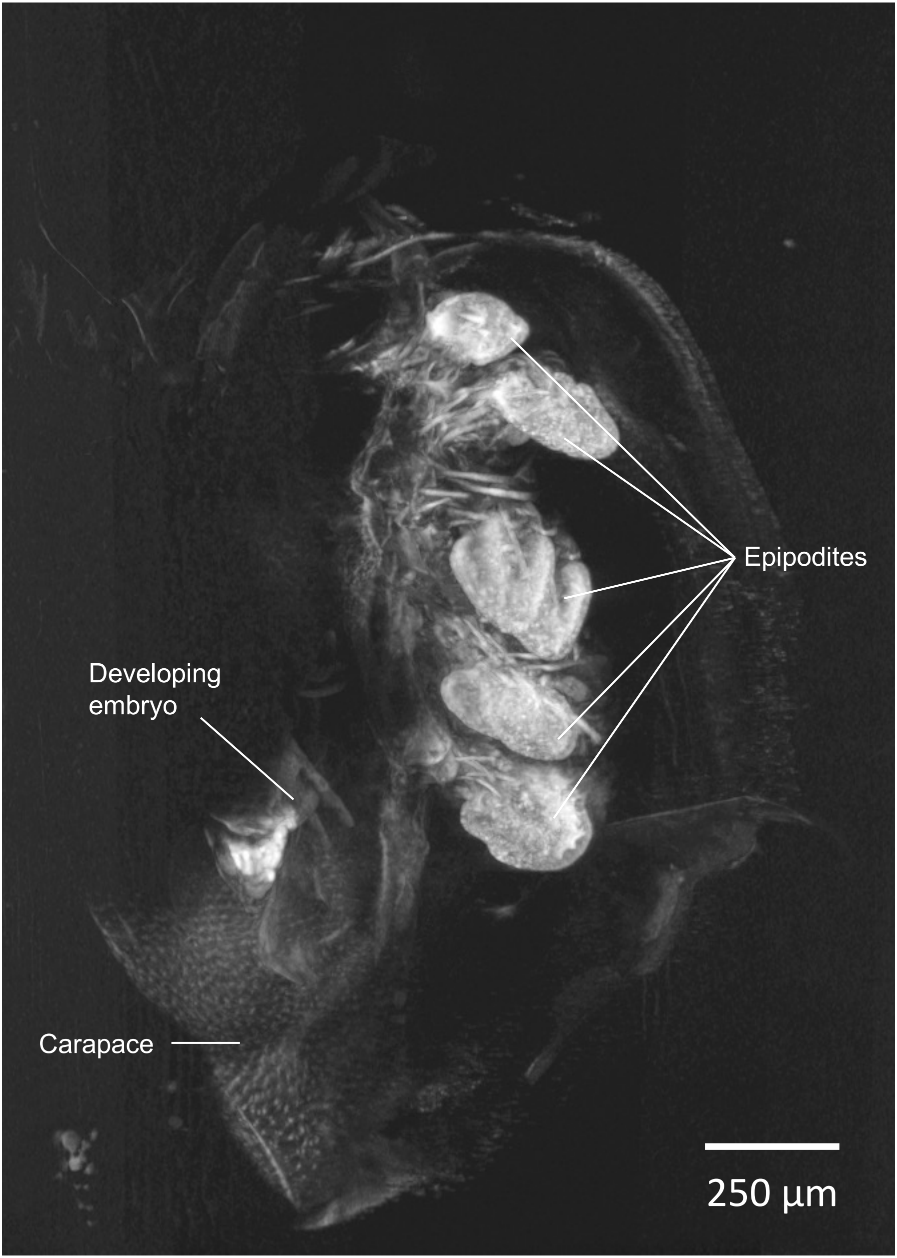

### S2 Fig

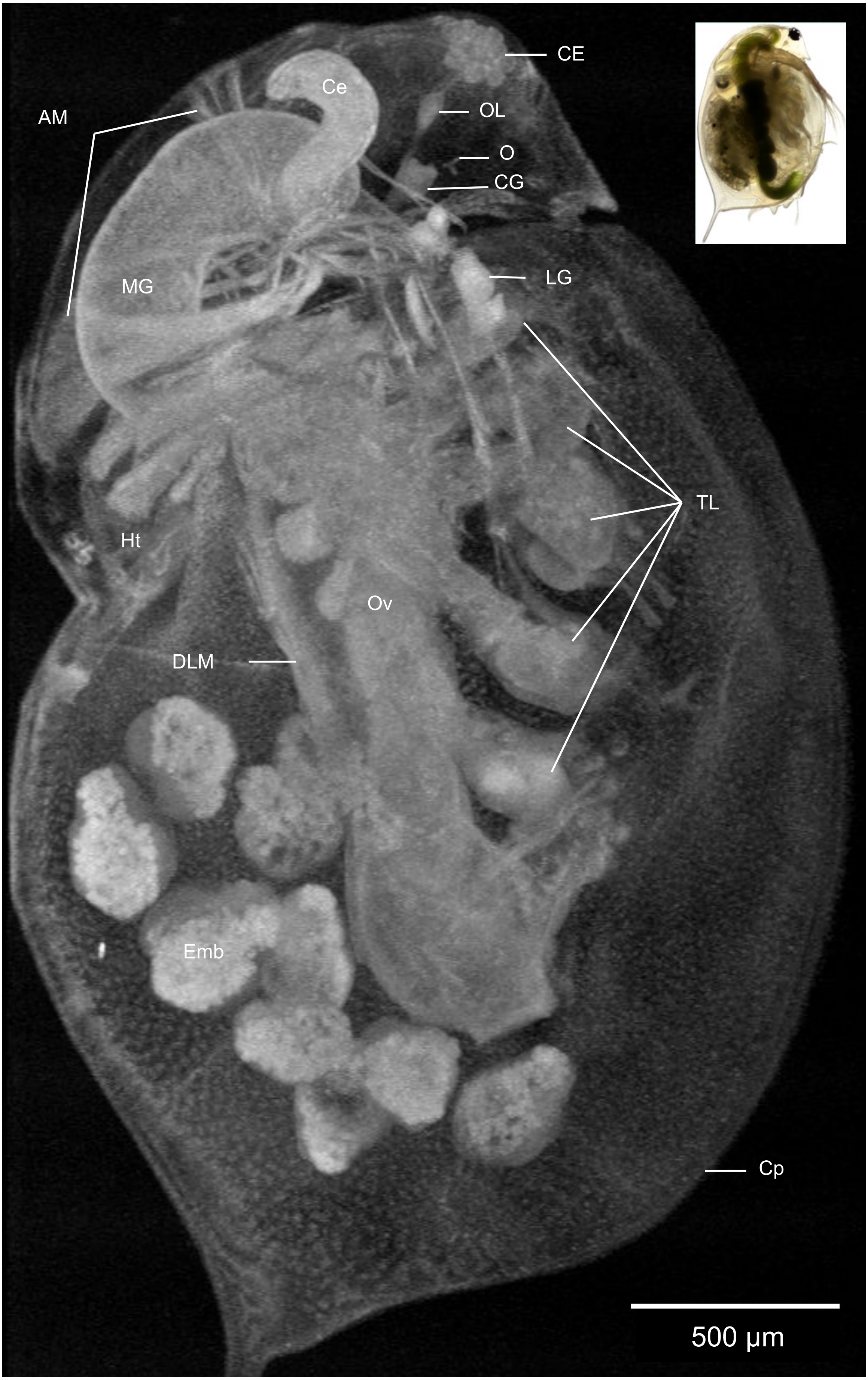
